# *Pseudomonas aeruginosa* synthesizes the autoinducers of its oxylipin-dependent quorum sensing system extracellularly

**DOI:** 10.1101/2021.09.08.459554

**Authors:** Eriel Martínez, Carlos J. Orihuela, Javier Campos-Gomez

## Abstract

The oxylipin-dependent quorum sensing system (ODS) of *Pseudomonas aeruginosa* relies on the production and sensing of two oxylipin autoinducers, 10S-hydroxy-(8*E*)-octadecenoic acid (10-HOME) and 7*S*,10*S* dihydroxy-(8*E*)-octadecenoic acid (7,10-DiHOME). Here, and contrary to the prevailing notion that bacterial autoinducers are synthesized intracellularly, we show that 10-HOME and 7,10-DiHOME biosynthesis occurs extracellularly, and this requires the secretion of the oxylipin synthases. We implemented a genetic screen of *P. aeruginosa* strain PAO1, which identified fourteen genes required for the synthesis of oxylipins. Among the identified genes, four encoded components of the ODS system and the other ten were part of the Xcp type II secretion system (T2SS). We created a deletion mutant of *xcpQ*, which encodes the outer membrane component of Xcp, and found it recapitulated the impaired functionality of the transposon mutants. Upon further examination, the lack of ODS function was demonstrated to be caused by the blocking of the DS enzymes secretion. Notably, the *xcpQ* mutant activated the ODS system when exposed to 10-HOME and 7,10-DiHOME, indicating that the sensing component of this quorum sensing system remains fully functional. In contrast with the detrimental effect previously described for T2SS in biofilm formation, here we observed that T2SS was required for robust *in vitro* and *in vivo* biofilm formation in an ODS dependent manner. To the best of our knowledge, this study is the first to find QS autoinducers that are synthetized in the extracellular space and provides new evidence for the role of the T2SS for biofilm formation in *P. aeruginosa*.

**IMPORTANCE:** We previously showed that the ODS quorum sensing system of *P. aeruginosa* produces and responds to oxylipins derived from host oleic acid by enhancing biofilm formation and virulence. Herein, we developed a genetic screen strategy to explore the molecular basis for oxylipins synthesis and detection. Unexpectedly, we found that the ODS autoinducer synthases cross the outer membrane using the Xcp Type 2 secretion system of *P. aeruginosa* and thus, the biosynthesis of oxylipins occur extracellularly. Biofilm formation, which was thought to be impaired as result of Xcp activity, was found to be enhanced as result of ODS activation. This is a unique QS system strategy and reveals a new way by which *P. aeruginosa* interacts with the host environment.

## INTRODUCTION

*Pseudomonas aeruginosa* is an opportunistic pathogen that can cause disease in plants, animals and humans with breaches in their mechanical or physiological defense barriers (1). One reason for this is that *P. aeruginosa* has a versatile battery of extracellular and surface-associated virulence factors and is therefore able to form recalcitrant biofilms that protect it from diverse conditions of stress, including the immune system and antibiotics (2). Quorum sensing (QS) is a bacterial cell-to-cell communication system that functions to regulates behavior at the cell community-level. QS systems have been shown to play a critical role in the regulation of virulence factors and biofilm formation in *P. aeruginosa* (3). Indeed, deletion of any of the previously described interconnected QS systems of *P. aeruginosa, las, rhl*, PQS and IQS, has been demonstrated to attenuate bacterial virulence in a variety of animal models (4). Recently we described a fifth QS system for *P. aeruginosa* that is regulated by oxylipins, hence named <u>O</u>xylipin-<u>D</u>ependent Quorum Sensing <u>S</u>ystem (ODS) (5). Similar to *P. aeruginosa*’s other QS systems, we showed that disruption of ODS strongly attenuated bacterial virulence in both plant and animal models (6). Accordingly, we showed that the ODS system regulates *P. aeruginosa* twitching, swarming, flagella-mediated swimming, which promotes biofilm formation.

Oxylipins are bioactive oxygenated lipids. In mammals, oxylipins are derived from polyunsaturated fatty acids by the action of cyclooxygenase, lipoxygenase, or cytochrome P450 oxygenase enzymes (7). They serve to modulate inflammatory pathways, but also have multiple other functions including antimicrobial properties. Oxylipins are also produced by invertebrates, plants, and fungi (8). Typically, oxylipins are not stored in tissues but are formed on demand from precursor fatty acids of endogenous sources (9). Pertinently, *P. aeruginosa* ODS relies on the presence and sensing of the extracellular oxylipins (10*S*)-hydroxy-(8*E*)-octadecenoic acid (10-HOME) and 7*S*,10*S*-dihydroxy-(8*E*)-octadecenoic acid (7,10-DiHOME)(10). We have shown these are synthesized by *P. aeruginosa* from exogenous oleic acid (OA) using the fatty acid diol synthase (DS) enzymes that are encoded by the DS operon (Fig 1A) (11).

**Figure 1.**
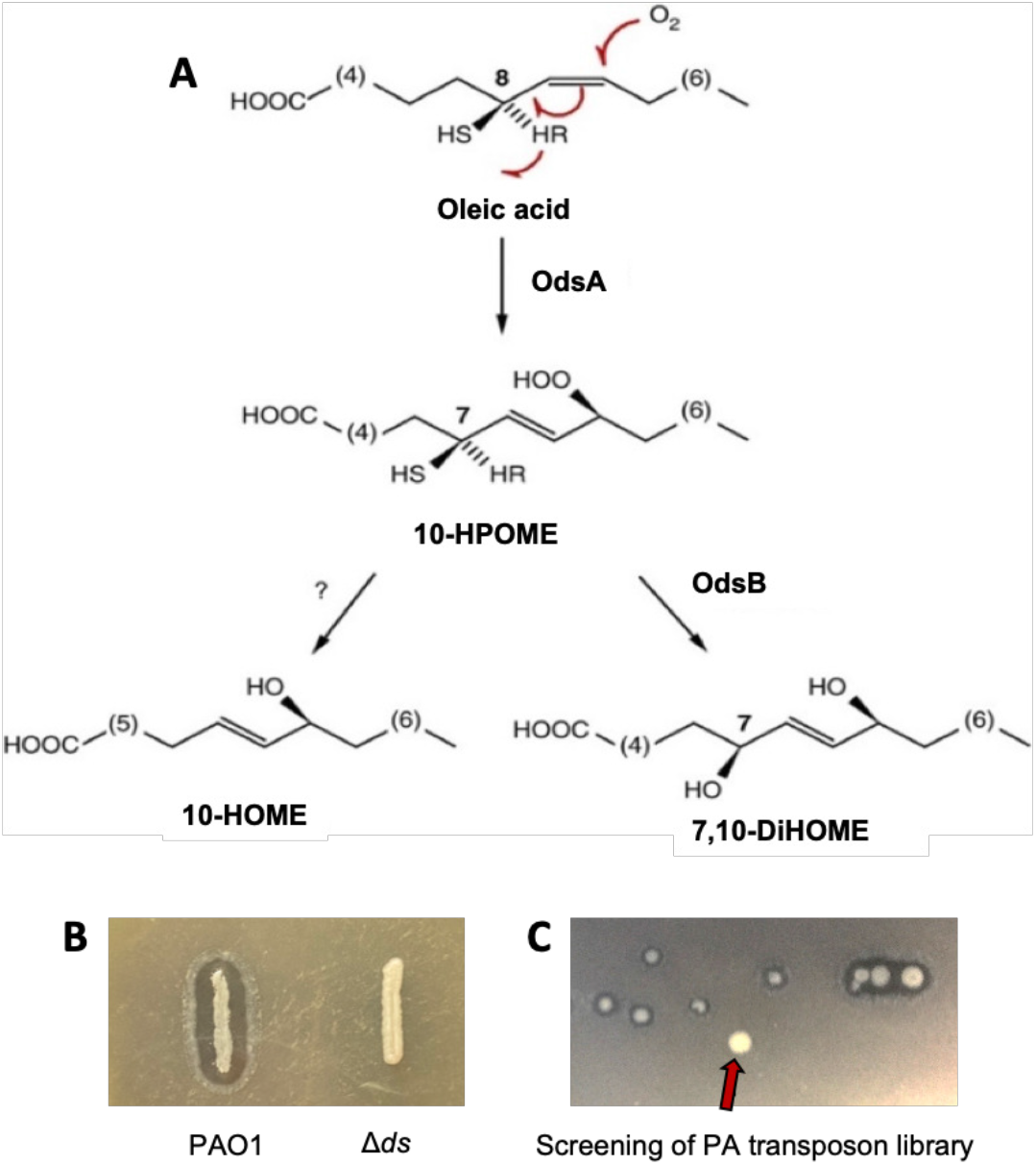
Screening assay for the search of genes affecting oxylipin production. **A)** Oxylipins biosynthetic pathway of *P. aeruginosa*. The enzyme 10(S)-dioxygenase (OdsA) transforms host oleic acid into 10S-Hydroperoxide-octadecenoic acid (10S-HPOME) by stereospecific oxygenation at position C10 of the oleic acid alky chain. Subsequently, 10S-HPOME could be isomerized by the enzyme (7*S*,10*S*)-hydroperoxide isomerase to form 7*S*,10*S*-DiHOME or be reduced to 10-HOME by an undefined mechanism. **B)** Picture showing the WT phenotype of *P. aeruginosa* vs that of the ΔDS deletion mutant when plated on LB agar + OA (1%). **C)** Representative picture of a plate section of our transposon screening showing a colony lacking the halo

Canonically, the biosynthesis of bacterial QS autoinducers occurs intracellularly using endogenous sources (12), and these are released to the extracellular space by a variety of means (13). The mechanisms by which the autoinducers reach the extracellular space depends on the nature of the autoinducers and the bacterial species; they include free diffusion through the bacterial membranes, the use of efflux pumps, and via outer membrane vesicles (14-16). Importantly, and prior to this report, we worked under the assumption that this held true for the ODS system, and that imported OA from the bacterial environment was converted within the periplasm of the bacteria to 10-HOME and 7,10-DiHOME by the DS enzymes (17). Herein we demonstrate that this is not the case, and that instead the DS enzymes are secreted from the periplasmic space via the Xcp type II secretion system (T2SS). Once outside the cell they use host-derived exogenous OA to synthesize the oxylipin autoinducers. To the best of our knowledge this is the first report of a QS system whose autoinducer molecules are synthetized in the extracellular space. They highlight the versatility of *P. aeruginosa* to sense and take advantage of the host environment.

## RESULTS

### Screening for *P. aeruginosa* factors involved in the production of the ODS autoinducers

The DS activity of *P. aeruginosa* introduces one or two hydroxyl groups into the alkyl chain of OA (Fig. 1A) (10, 18); the oxylipins derived from this activity being more hydrophilic than the OA substrate. In addition, these oxylipins have surfactant emulsifying properties that help to dissolve OA in suspension above the critical micelle concentration. We noticed that these properties of oxylipins enable wildtype *P. aeruginosa* colonies to produce a transparent halo when this bacterium is grown on Lysogeny Broth (LB) agar plates containing 1% OA, which renders the medium opaque. Consequently, *P. aeruginosa* lacking DS activity, i.e. the ΔDS mutant, do not from this halo when grown under the same conditions (Fig. 1B).

We took advantage of this characteristic phenotype to identify the bacterial genes involved in oxylipin production. To do this we performed a genetic screen using a sequence-defined transposon (*ISphoA/hah* or *ISlacZ/hah*) insertion library created in the model strain PAO1 (19). In total we screened more than 30,000 independent clones for the inability to form the halo on LB-agar supplemented with 1% of OA (Figure 1C). We identified 31 colonies unable to form the transparent halo even after re-streaking on fresh plates. We successfully PCR amplified and sequenced the DNA regions flanking the transposon for all the 31 mutants, thereby identifying 15 genes putatively required for oxylipin production (Table 1).

**Table 1:**
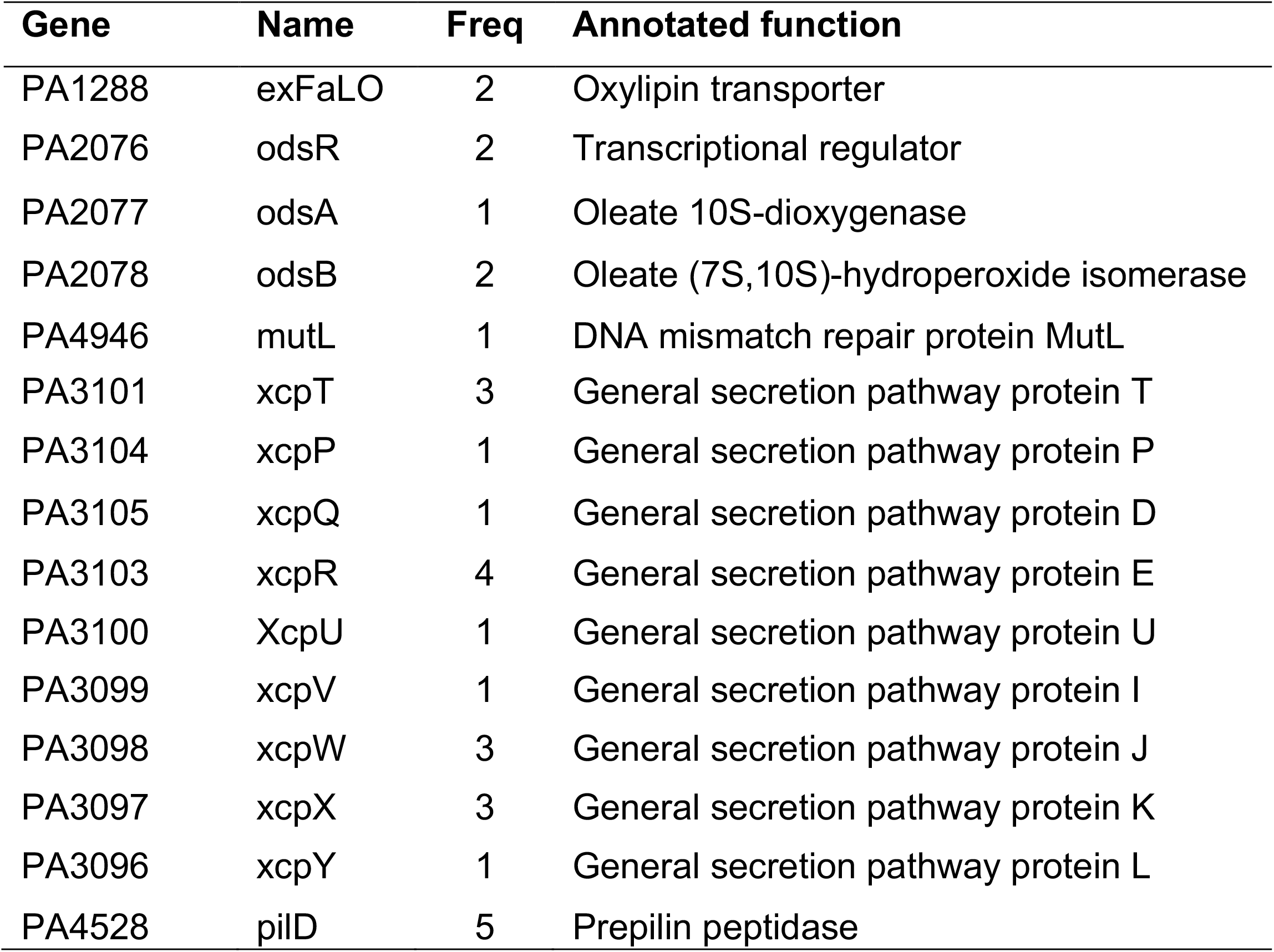
Genes identified in the screening, their frequencies and functions.

Analysis of the identified clones revealed that three of them carried a transposon insertion in the DS operon, which encodes the DS enzymes OdsA and OdsB that catalyze oxylipin biosynthesis (annotated as PA2077 and PA2078, respectively, in the pseudomonas database). Two other transpositions occurred in *odsR* (PA2076), the DS operon transcriptional regulator found immediately upstream and in opposite orientation to the DS operon. Additionally, two transpositions were identified in the outer membrane transporter encoded by PA1288, *exFadLO*, which we previously had proposed to be involved in oxylipin export across the outer membrane (17). Based on our previous knowledge on the components required for oxylipin biosynthesis, the above-mentioned transposition events were expected to be found, and thereby confirmed the validity of the screening strategy. Transposon insertions providing new insight into the ODS system included nineteen transpositions in genes encoding 9 distinct components of the Xcp T2SS; all of which are found in the Xcp region of PAO1 chromosome (PA3095 to PA3105). Another five transpositions occurred in *pilD*, the gene encoding a prepilin peptidase, which is vital for the processing of some T2SS components in *P. aeruginosa* (20). Lastly, one transposition occurred in *mutL*, which encodes a DNA mismatch repair enzyme which promotes large chromosomal deletions in *P. aeruginosa* (21). Subsequent studies using a clean deletion *mutL* mutant failed to corroborate the halo deficient phenotype, suggesting that the colony phenotype of the *mutL* transposition found, was most likely due to a secondary mutation caused by the consequent MutL deficient hyper-mutagenic phenotype.

### Oxylipin synthases cross the outer membrane via Xcp T2SS

T2SS are involved in the transport of proteins across the outer membrane from the periplasmic space (22). In *P. aeruginosa* Xcp has been demonstrated to be involved in the translocation of multiple proteins including established virulence factors (23). To confirm the role of the T2SS in oxylipin production we made an in-frame deletion of the gene *xcpQ* (Δ*xcpQ)*, which encodes the outer membrane porin component of the T2SS. As expected, colonies of Δ*xcpQ* failed to produce transparent halos when plated on LB containing 1% of OA (Fig 2A). In addition, using thin layer chromatography, we confirmed that Δ*xcpQ* failed to produce any 10-HOME or 7,10-DiHOME in the extracellular space when grown in LB supplemented with OA (Fig. 2B). Importantly, Δ*xcpQ* deficient mutant complemented *in trans* with a wild type *xcpQ* gene restored the halo and production of oxylipins (Fig 2A, B). These results indicate that the DS enzymes most likely cross the outer membrane using the T2SS. In agreement with this notion, we found that when PAO1 and Δ*xcpQ* were treated with exogenous 7,10-DiHOME to induce DS gene expression, the DS enzymes accumulated into the periplasm of the mutant, but not in that of the wildtype (Fig. 2C). Moreover, contrary to the wildtype controls, cell free supernatants of Δ*xcpQ* did not show DS activity (Fig. 2D).

**Figure 2.**
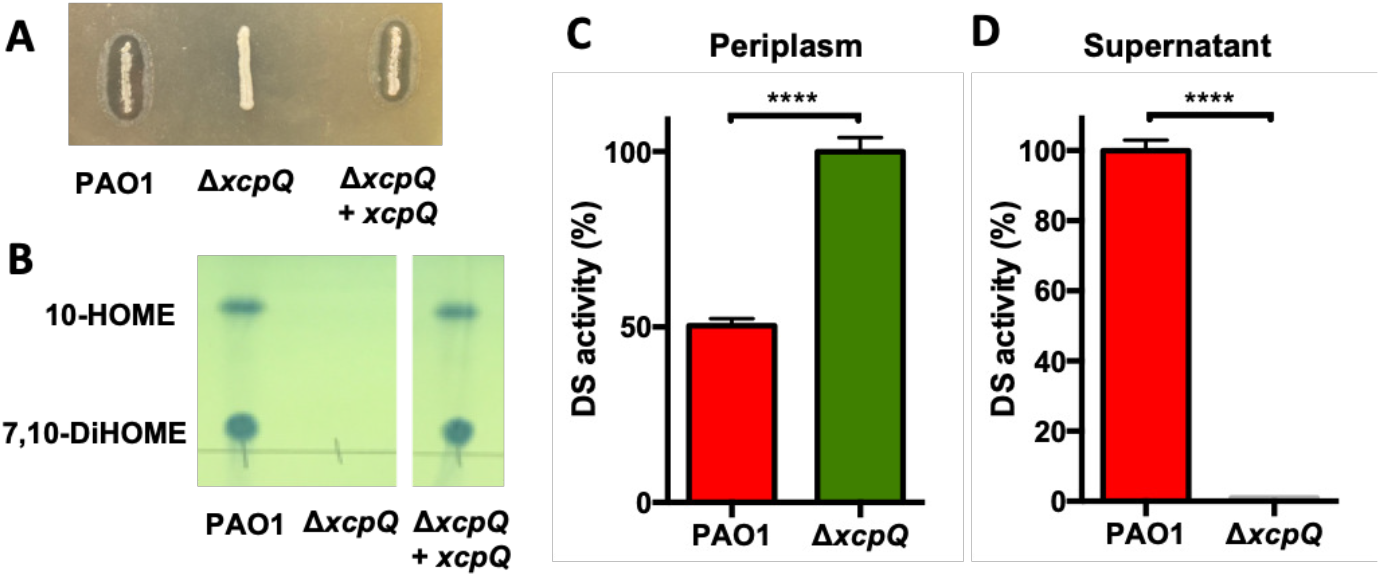
Accumulation of 7,10 DiHOME oxylipin in culture supernatants of PAO1 and its isogenic mutant Δ*xcpQ*. **A)** The Δ*xcpQ* mutant failed to produce 7,10-DiHOME in the culture supernatant. The production of 7,10-DiHOME was restored by Δ*xcpQ* when it was complemented with a wild type *xcpQ gene* expressed from a plasmid. **B)** Think layer chromatography (TLC) analysis of oxylipins 10-HOME and 7,10-DiHOME accumulated in the supernatant of PAO1 and Δ*xcpQ* complemented in trans. **C)** The DS enzymes accumulate in the periplasm of Δ*xcpQ*. **D)** The Δ*xcpQ* mutant shows a negligible DS activity in the culture supernatant.

### ODS functionality requires DS enzymes secretion through the T2SS

The ODS system involves accumulation of oxylipins in the extracellular medium and subsequently sensing of the oxylipin signal (5). Accumulated oxylipins induce the expression of several genes in *P. aeruginosa*, previously identified by our group (5). To determine the impact of the T2SS in the expression of genes under the control of ODS, we followed the expression kinetic of a representative ODS regulated gene, PA3427 (5). For this, we made a genetic fusion of PA3427 with the *lacZ* reporter gene. As expected, in a Δ*xcpQ* background PA3427-lacZ was unresponsive to the presence of OA (Fig. 4A). To discard any collateral effect that a dysfunctional T2SS might have on the expression of PA3427 we corroborated that this gene was expressed normally in Δ*xcpQ* at the same level of PAO1 when induced with the purified oxylipin 7,10-DiHOME (Fig. 4B). Thus, Xcp is required for DS secretion leading to extracellular oxylipin production, however, it is not involved in the sensing or response to the oxylipin signal.

**Figure 3.**
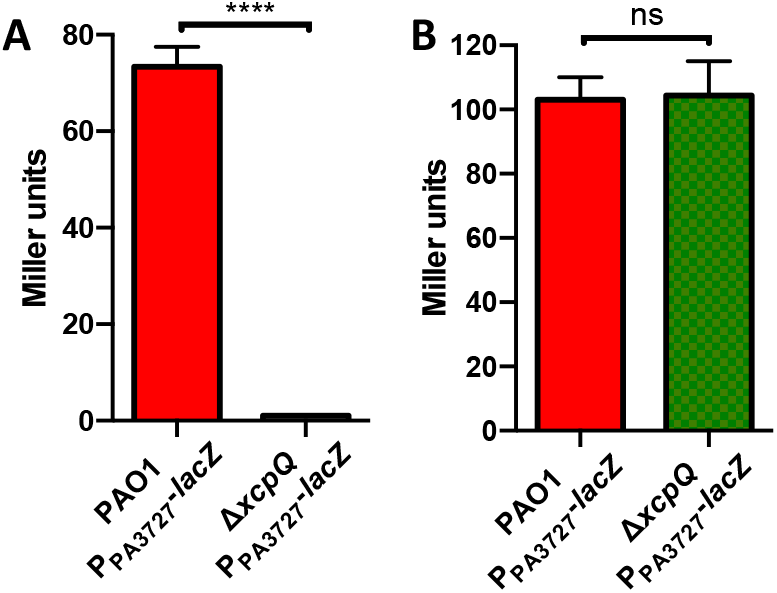
Beta-galactosidase activity of P_PA3427_-lacZ fusion in PAO1 and Δ*xcpQ* genetic backgrounds. **A)** Δ*xcpQ* mutant showed a negligible PA3727-lacZ expression in the presence of OA. **B)** Δ*xcpQ* mutant showed the same level of expression as PAO1 when induced with 7,10-DiHOME.

**Figure 4.**
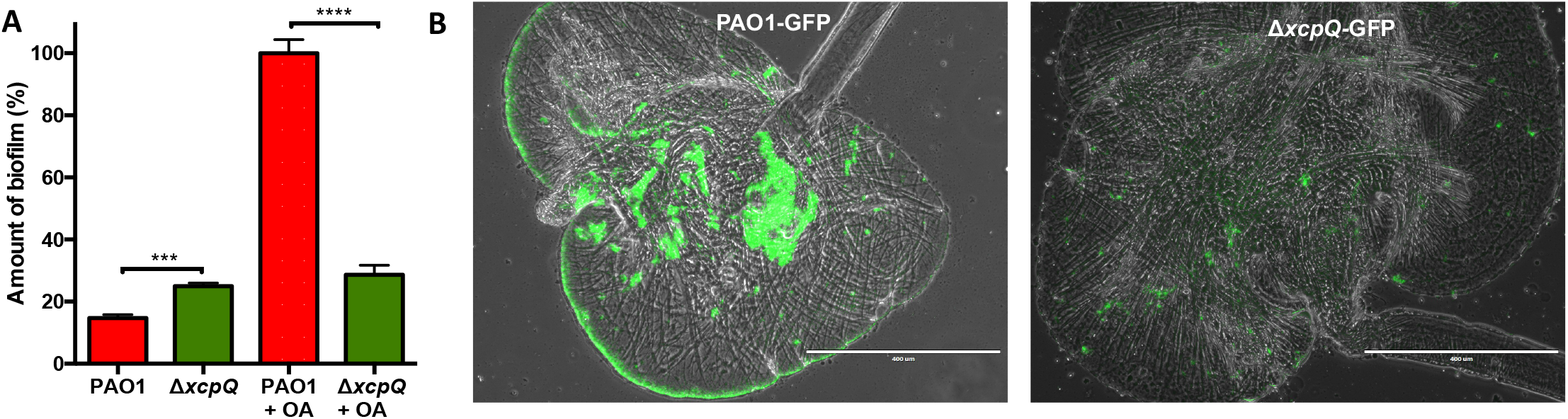
Biofilm formation by PAO1 and its isogenic mutant Δ*xcpQ*. **A)** Δ*xcpQ* produces higher amount of biofilm than WT PAO1 when tested in the absence of OA. In contrast, it produces more biofilm than WT in media supplemented with OA. **B)** Δ*xcpQ* produced less amount of biofilm in media supplemented with oleic acid. C) Fluorescence microscopy analysis of crops from *Drosophila melanogaster* infected with PAO1 expressing GFP (PAO-GFP) or Δ*xcpQ* expressing GFP (Δ*xcpQ-*GFP). PAO1 formed more biofilm that Δ*xcpQ*. Bars represent 400 μm. The size/ resolution for each panel was adjusted to 2.125 1.587 in/600 dpi from 17.770 13.333 in/72 dpi of the originals. Pictures are representative of three independent experiments.

### The T2SS is linked to biofilm formation in *P. aeruginosa*

The Xcp system of *P. aeruginosa* promote biofilm dispersal. Consequently, the level of Xcp secretome and biofilm formation under static conditions have an inverse correlation (24). In agreement with this prior report, we observed an increase in the amount of biofilm formed by Δ*xcpQ* compared to the WT PAO1 strain using the microtiter plate model, when both were grown in the absence of OA (Fig. 5A). However, in the presence of OA the amount of biofilm formed by PAO1 was significantly higher than that of Δ*xcpQ* (Fig. 5A). This result agreed with our previous study reporting that ODS system promotes biofilm formation *in vitro* (6). We also recapitulated this result *in vivo* using *Drosophila melanogaster* fed with oleic acid-supplemented food. Imaging of fly crops conclusively showed reduced biofilm formation for Δ*xcpQ* versus the PAO1 control (Fig. 5B). Thus, the T2SS of *P. aeruginosa* promote biofilm formation provided that OA is available and a functional ODS system is present.

**Figure 5.**
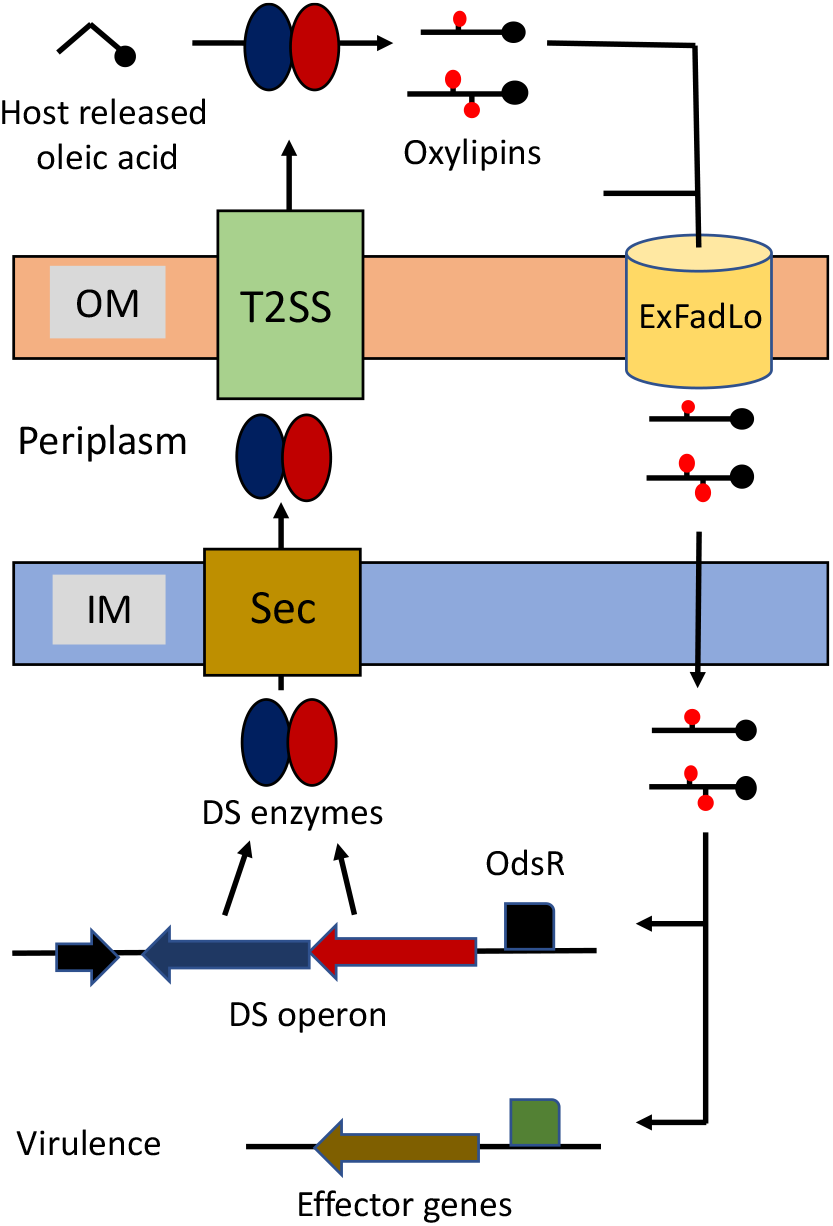
Proposed model of ODS. At low cell density the DS operon is weakly expressed. When the cell density increases and host oleic acid is present, the DS enzymes start to produce oxylipins. Subsequently the oxylipins enter the cells through ExFadLO and bind OdsR which in turn induces the DS operon. Induced DS enzymes cross the inner membrane through the Sec secretory pathway to reach the periplasm. Then the enzymes cross the outer membrane via Xcp type 2 secretion system (T2SS). This causes a sudden accumulation of extracellular DS enzymes and oxylipins that activate the ODS system and induce the effector genes.

## DISCUSSION

Herein we report the first example of extracellularly produced QS autoinducers. The autoinducer synthases OdsA and OdsB of the *P. aeruginosa* ODS system are exported to the extracellular space through the Xcp T2SS. Once in the extracellular space, the DS enzymes use exogenous OA as a substrate to synthetize the oxylipin inducers 10-HOME and 7,10-diHOME, which in turn enter into the bacterial cells, presumably via the ExfadLO transporter (See proposed model in Fig. 6). In addition, this study establishes a new link between the T2SS, QS and biofilm formation in the relevant opportunistic pathogen *P. aeruginosa*.

In *P. aeruginosa*, Xcp is involved in the secretion of several virulence factors, such as the elastase LasB, the lipase LipA and the alkaline phosphatase PhoA (25). *P. aeruginosa* has a second T2SS known as Hxc (26), but this is thought to have a considerably restricted role and we observed no evidence of it being involved with the ODS system. Our finding that the Xcp also exports OdsA and OdsB, increases the repertoire of virulence factors known to be secreted by this general secretory pathway. All described T2SS in Gram-negative bacteria employ a two-step process to secrete proteins from the cytoplasm to the extracellular space through a transient periplasmic intermediate. The first step of translocation through the inner membrane is commonly carried out by the Sec or Tat systems (27, 28). Analysis of the amino acid sequence identified a putative N-terminal signal peptide in both OdsA and OdsB suggesting both use the Sec secretory pathway for translocation through the inner membrane, although this remains to be demonstrated (29). Noticeable, the fact that we did not detect transposition events in the genes encoding components of the Sec secretory pathway is likely due to the essential role for viability of this system in most bacteria (30, 31).

In our genetic screen we identified two independent transposition events in *exfadLO*, which confirms that the encoded transporter is essential for the normal functioning of the ODS system. Prior to this report, we had postulated that ExFadLO was responsible for the export of intracellularly synthesized oxylipins. In light of our finding that oxylipins are synthesized extracellularly as result of Xcp-dependent secretion of DS enzymes, we now instead propose that one of its main functions is to import oxylipins. Ongoing studies in the laboratory are testing this hypothesis.

Biofilm formation is an important mechanism by which bacteria establish themselves within a host as well as a mechanism of defense from host factors. OA is an abundant fatty acid in host tissues, and we previously demonstrated that *P. aeruginosa* scavenges OA from the host to produce oxylipins, which promotes biofilm formation *in vivo* (6). Here we show that the Xcp T2SS promotes biofilm formation, both *in vitro* and *in vivo*, in the presence of OA in an ODS-dependent fashion. The convergence of these traits makes sense as OA is a host derived signal, specially from wounded ones, and formation of biofilm as a result of ODS activation would be a mean to adapt to the host environment.

In summary, we developed a simple screening strategy that allowed the identification of bacterial genes required for ODS functioning in *P. aeruginosa*. As a result, we found that the Xcp T2SS of *P. aeruginosa* translocate the DS enzymes, which synthesize the ODS autoinducers, through the outer membrane. This was an unexpected finding since the biosynthesis of previously known QS autoinducers occur intracellularly and subsequently released into the extracellular space. ODS is an environment specific QS system, which depends on the presence of exogenous OA. Thus, we propose that the translocation of the DS enzymes to the extracellular media, which is peculiar among autoinducer synthases, is a way for this QS to be fine-tuned to the host environment, allowing *P. aeruginosa* to respond quickly to this condition.

## MATERIALS AND METHODS

### Bacterial strains, plasmids and culture conditions

Strains, plasmids and oligonucleotides used in this study are described in Table 2. Used strains were routinely grown in lysogeny broth (LB) medium at 37°C, to which agar was added when solid medium was required. When required, *P. aeruginosa* was grown in M63 media supplemented with 0.2% glucose, 0.1% casaminoacids and MgSO_4_ 1 mM (M63 complete). Antibiotics were added, when necessary, at the following concentrations: Ampicillin (Amp), 100 μg/mL; Carbenicillin (Cb), 300 μg/mL (*P. aeruginosa*); Chloramphenicol (Cm), 25 μg/mL for *Escherichia coli* and 200 μg/mL for *P*.*aeruginosa;* Kanamycin (Km), 25 μg/mL. OA 90% (Sigma) was added to cultures for oxylipin production and purification. M63 complete or LB media were supplemented with OA 99% (Sigma) or purified oxylipins at the specified concentrations when required. LB agar without NaCl plus 15% sucrose was used to segregate suicide plasmids from merodiploids during construction of *xcpQ* deletion mutant (Δ*xcpQ*) strain by allelic exchange (see below).

**Table 2:**
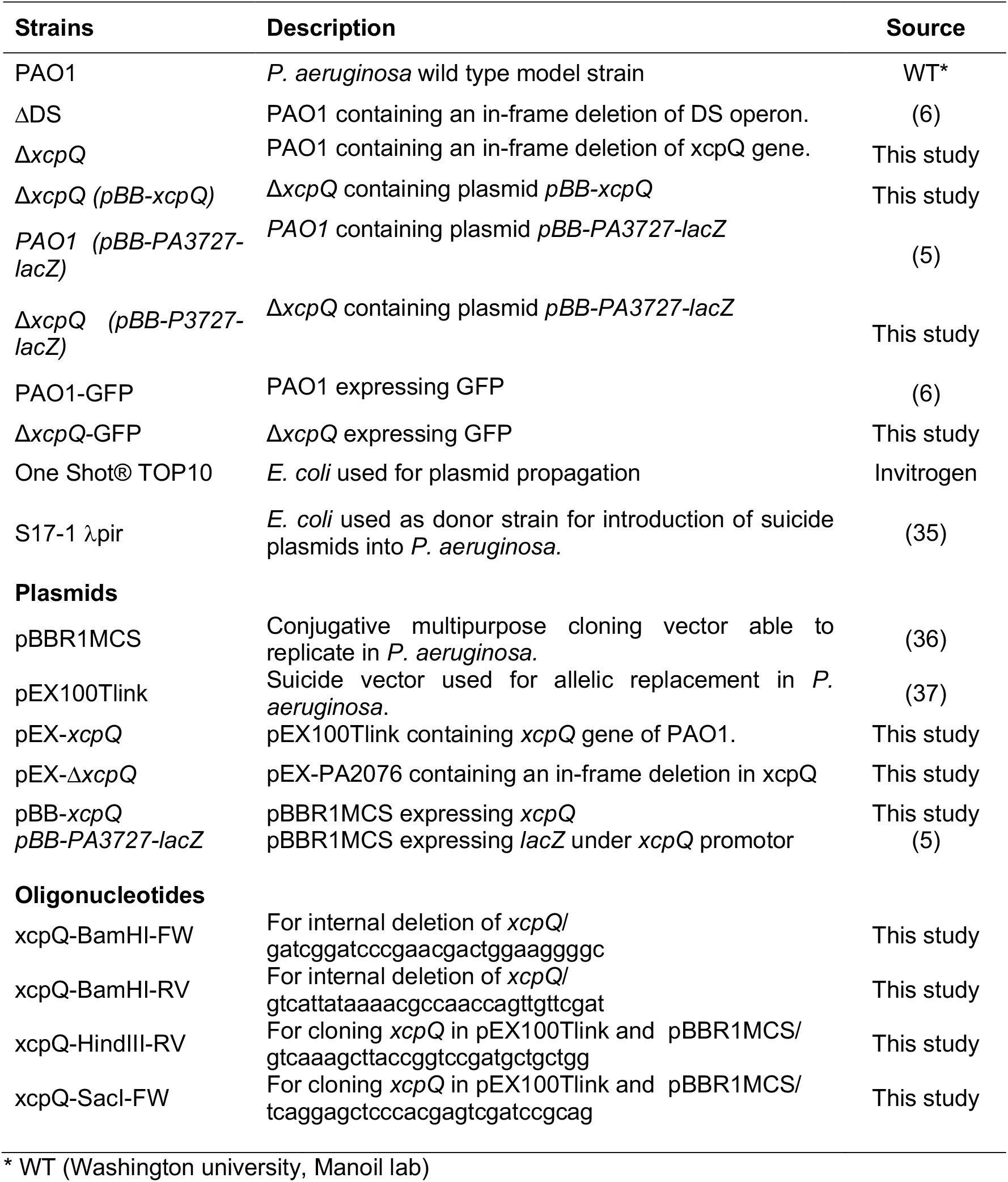
The list of strains, plasmids and oligonucleotides used in this study.

### Transposon library screening and mutant characterization

A random mariner transposon library of *P. aeruginosa* was acquired from the University of Washington (ref). The transposon library was amplified by growing the bacteria in LB broth up to OD_600_ = 1 (exponential phase) and proper dilutions of the bacterial suspension were plated on LB agar supplemented with 1 % OA to obtain separate clones. More than 30,000 independent colonies were obtained and from those, the clones lacking a transparent halo were selected for further analysis. The gene mutation of each selected clone was identified by DNA sequencing as previously described (19).

### Genetic constructions

A region of *P. aeruginosa* PAO1 genome comprising the *xcpQ* gene and its flanking regions (∼550 bp of each flank) was amplified using primers *xcpQ*-F-*Sac*I and *xcpQ*-R-*Hin*dIII (Table 1). The primers introduced *Sac*I and *Hin*dIII restriction sites at the extremes of the amplified fragment, which were used to insert the fragment into the pEX100Tlink suicide vector^2^ digested with the same enzymes to obtain pEX-*xcpQ* plasmid. Subsequently, an internal fragment of 1,867 bp was deleted from the *xcpQ* gene by doing a reverse PCR amplification using pEX-*xcpQ* as template and primers Δ*xcpQ*-F-*Bam*HI and ΔxcpQ-R-*Bam*HI. The amplified fragment was digested with *Bam*HI and self-ligated. The obtained plasmid, which was named pEX-Δ*xcpQ*, contains *xcpQ* gene with an internal in-frame deletion flanked by approximately 550 bp by each side (required for homologous recombination). This suicide plasmid was used to delete *xcpQ* from PAO1 chromosome by allelic replacement. Genetic fusion of PA3427 with the reporter gene *lacZ* was done as previously described (5).

### Periplasm extraction

The periplasm was extracted following the method of Wood (32) with modifications by Robles-Price *et al*. (33). Briefly, cells were collected by centrifugation (4,000 × g, 10 min, 4°C), washed twice in 30 mM Tris-HCl, 150 mM NaCl (pH 7.1), and kept in ice for no longer than 1 h. Periplasm was further obtained by suspending the cells in 6 mL of 30 mM Tris-HCl, 20% sucrose, 4 mM EDTA, 0.5 mg/mL lysozyme, 1 mM PMSF (pH 8), and subsequent 60 min incubation at 30°C with gentle shaking. MgCl_2_ was added at 10 mM final concentration as soon as the suspension reached 30°C. Finally, the suspension was centrifuged (11,000 × g, 15 min, 4°C), and the supernatant containing the periplasmic fraction was collected.

### β-Galactosidase activity assay

*P. aeruginosa* strains to be assayed were grown overnight in LB agar plates, then bacterial suspensions were prepared in fresh M63 to OD_600_ = 0.5 with or without oxylipins or OA (0.1 mg/ml). Cultures were incubated at 30°C for two hours and then 250 µl of each culture was mixed with 250 µl of Z buffer [Na_2_HPO_4_.7H_2_O (0.06M), NaH_2_PO_4_.H_2_O (0.04M), KCl (0.01M), MgSO_4_ (0.001M), *β* - mercaptoethanol (0.05M), pH to 7.0], 50 µl of 0.1% SDS and 100 µl of chloroform and the mix vortexed for 20 sec. The tubes were incubated at 30°C for 5 min and the reaction started by adding 100 µl of *o*-nitrophenyl-β-D-galactoside (ONPG, 4 mg/ml) and briefly vortex mixing. Reactions were incubated at 30°C for 1 hour and stopped by adding 250 µl of 1M Na_2_CO_3_. The OD at 420 nm and at 550 nm was measured for each tube. Finally, β-Gal activity was calculated using the equation: Miller Units = 1,000 x [(OD_420_ - 1.75 x OD_550_)] / (T x V x OD_600_); where OD_420_ and OD_550_ are the final reads from the reaction mixture, OD_600_ is the initial cell density of the cultures, T is the time of the reaction in minutes, and V the volume of culture used in the assay in mL.

### Thin layer chromatography (TLC)

TLCs were run on 60 Å silica gel plates of 20 × 10 cm and 200 µm thickness (Whatman^®^). The mobile phase solvent was a mix of hexane, ether and acetic acid (80/20/5). TLC plates were revealed with 10% phosphomolybdic acid in ethanol. The relative amount of oxylipins were semi-quantitated by densitometry of TLC spots using ImageJ software.

### Purification of 7,10-DiHOME oxylipin

7,10 Di-HOME was purified as previously described (6). Briefly, PAO1 was plated in LB agar and incubated overnight at 30°C. The bacterial biomass was scraped from the plate and used to inoculate 200 ml of M63 complete supplemented with 1% OA. The culture was allowed to produce oxylipins and then centrifuged at 8000 x g for 15 min to remove bacterial cells. The supernatant was recovered and acidified (pH=2) with acetic acid glacial. Then a 1 vol/vol organic extraction with ethyl acetate was carried out and the organic phase was evaporated. The dried mixture obtained was dissolved in 3 mL of ethyl acetate and used for purification of 7,10-DiHOME using an Isco Teledyne Combiflash Rf 200 with four channels with 340CF ELSD (evaporative light scattering detector). Universal RediSep solid sample loading pre-packed cartridges (5.0 g silica) were used to absorb the crude product and purified on 24 g silica RediSep Rf Gold Silica (20–40 µm spherical silica) columns using an increasing gradient of ethyl acetate (solvent B) over hexane (solvent A). Fractions collected for each detected peak were combined and evaporated, then dissolved in ethanol. The purity of the 7,10-DiHOME was checked by HPLC/MS analysis as previously described (10).

### Biofilm assay inside *Drosophila* crops

*P. aeruginosa* colonization of *D. melanogaster* crop was performed as previously described by Mulcahy *et al* (34). All experiments were performed with 3-day-old *D. melanogaster* from both sexes of the WT Oregon R (acquired from Carolina Biologicals Company). *P. aeruginosa* strains constitutively expressing GFP were cultured on LB agar plates. Bacteria were resuspended in LB to OD600 = 1. Then 100 µl of the suspension was spotted onto a sterile filter (Whatman) that was placed on the surface of 5 mL of LB agar supplemented with 5% sucrose and 1% oleic acid. Flies were allowed to grow under this condition for 20 hours and then killed. Crops were placed on a drop of PBS on a microscope slide, sealed with a coverslip and observed using an EVOS FL Cell Imaging System. Pictures were captured using the same settings for each picture.

### Statistical analysis

Data are representative of three technical replicates and three biological replicates of each condition. Means plotted and a Student’s unpaired *t*-test (two-tailed) was used to determine differences between means of varying conditions after it was determined that the variance was similar between groups. All statistical analyses were performed using GraphPad Prism 8.3.1 software.

## Data availability

The authors declare that the data supporting the findings of this study are available within the article and its supplementary information files, or from the corresponding author upon request.

